# A Phylogeny-Based Approach to Discover Mutational Processes in Primates

**DOI:** 10.1101/2024.12.02.626204

**Authors:** James C. Kitch, Vladimir Seplyarskiy

## Abstract

The accumulation of germline mutations underpins population diversity and drives genetic evolution. Despite the availability of extensive phylogenetic data, the lack of suitable methodologies has hindered the comprehensive characterization of germline mutational processes across evolutionary trees. To address this, we develop a robust three-step methodology that extracts germline mutational processes from alignments of closely related species. First, we estimate regional, branch-specific trinucleotide mutational spectra from a multispecies alignment. Second, we extract mutational processes jointly across an evolutionary clade by analyzing mutation rate variation along the genome using Reciprocal Principal Components Analysis (RPCA). Finally we filter artifactual mutational signatures using DNA symmetry. Applying this method to five primate clades and a rodent outgroup clade revealed nine distinct mutational processes. Notably, five of these processes were consistently observed across all six groups. We identified underling biological mechanism for at least seven of the processes, highlighting phenomena such as biased gene conversion, bulky lesion resolution, and maternal mutagenesis. We validated identified processes using human and non-human polymorphism data. This study offers new insights into the biology and evolution of mutagenesis in primates and introduces a methodological toolkit to investigate mutational processes across phylogenies.

## 1 Introduction

Mutagenesis in germline produces variation that can be passed through generations. It is naturally of interest across multiple biological disciplines, including mendelian genetics ^1^, population genomics ^2,3^, phylogenetics ^4^ and finally evolutionary ^5,6^ and molecular biology ^7,8^. One of the most common approaches to interpret mutational patterns in germline is based on comparison with known cancer signatures ^9–12^, highlighting better understanding of somatic mutagenesis. Alternatively, it is possible to utilize unique features of germline mutagenesis, like differences in paternal and maternal gametogenesis ^13,14^, effect of meiotic recombination ^15,16^ or comparison of mutational patterns between populations ^17,18^.

Germline mutations are responsible for any standing genomic variation, and indeed it is possible to study them from different data modalities. Direct sequencing of sperm cells ^11,19,20^, family sequencing and extraction of de novo mutations ^21^, segregating variation in human population ^2^ and interspecies divergence ^22^ all allow insight into patterns of germline mutation. Recently, a massive dataset of rare human polymorphisms provided an opportunity to agnostically extract a comprehensive set of biologically interpretable mutational processes ^23^. However, this approach offered a static picture of germline mutagenesis in humans and does not provide insights into its evolution. Indeed, it has been shown that processes governing the accumulation of DNA changes evolved in recent human history ^18,24^ and is quite variable on the level of apes ^25,26^, mammals ^22^, and vertebrates ^5^. The observed strong negative association between life span and yearly somatic mutation rate ^27^ further enhances interest in understanding the dynamics of mutation rate evolution.

In this study, we leverage recently expanded phylogenetic data on primates ^28^ to extract spatial mutational processes from divergence data. To this end, we develop a procedure to calculate local mutation rates, based on single base substitutions, while accounting for trinucleotide context. We subsequently analyze matrices of spatial mutational patterns for multiple closely related species. Overall, we detected just nine mutational processes across six clades, with each of these processes being present in at least two clades.

We believe that both methods and resource that we created have very high utility and could be used to study dynamics of mutagenesis at unprecedented level and some of the signatures could be used as a close proxy, to DNA features, like spatial recombination rate, which is unknown for the most of studied species.

## 2 Data and Methods

### 2.1 The Progressive Cactus Alignment

The Zoonomia Project’s Progressive Cactus Genome Alignment^29^ is the foundation for our analysis of branch-specific mutational processes. This alignment, which contains aligned whole genome sequences for 241 mammalian species, allows inferences of species-specific mutation rates without the large sample size requirements of polymorphism data. However, the chromosomal duplications, rearrangements, and other large-scale sequence changes that happen between two species’ genomes during speciation makes phylogenetic data particularly vulnerable to alignment errors ^30^. In the face of this data quality challenge, we focus on inferring mutational processes on a smaller, forty-species subset of the tree (Figure 1). This subset mostly consists of primate genomes with one outgroup of hamster-related genomes. Primate genomes tended to be the highest quality genomes in the overall alignment, and short branches between species within a subclade decrease the prevalence of chromosomal rearrangements. Even on these higher-quality alignments, extracting mutational signatures is ripe with challenges. The next section details our data-cleaning strategies for mitigating the noise in these alignments.

**Figure 1:**
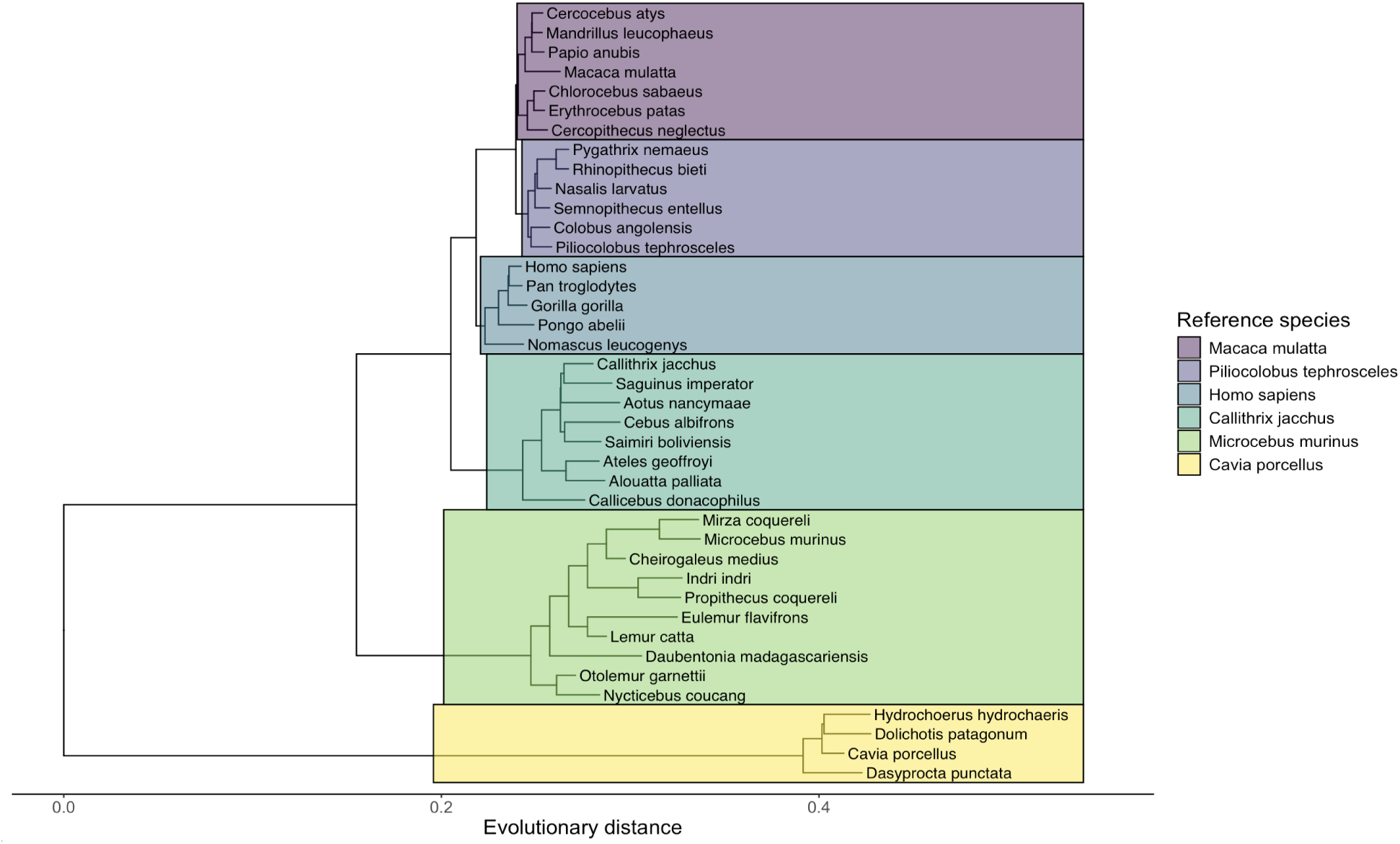
Phylogenetic tree of all species included in this analysis. Colors refer to evolutionary clades and are labeled by their assigned reference species.

#### 2.1.1 Reference species selection

We accessed the Cactus alignment in its native, reference-free Hierarchical Alignment (HAL) file ^31^ format. The HAL file format is optimized for sequence storage and flexibility, but attempting any large-scale analysis requires assigning reference species and working with Multiple Alignment Files (MAF). We group species based on distinct evolutionary clades (see Figure 1), and assign reference species based on scaffold N50 as an approximate metric of sequence quality ^32^.

#### 2.1.2 Data filtering and cleaning

After assigning reference species, we implement three additional filtering steps to increase the quality of the final alignment.

First, we keep only one species from any set of species that share the same genus (e.g. *Pan paniscus* and *Pan troglodytes*). In these cases, we keep the species with the higher sequence quality as measured by scaffold N50. Whereas long terminal branches are undesirable because they increase the likelihood of alignment errors, excessively short branches are also problematic for multiple reasons. First, short branches do not allow for a sufficient number of mutations to accumulate in order to accurately study mutational spectra. Second, incomplete linage sorting (ILS)–or inconsistencies between the gene geneaology and the species tree– is especially problematic on short branches because of comparable roles of ancestral polymorphisms and divergence ^33^. The variation of topology along the genome caused by ILS would affect the local variability of mutation rate and artificially impact inferred mutational processes.

Next, we remove all sequences (e.g. contigs) within an alignment with total length less than 10MB, as shorter sequences suggest fragmented, low-quality sections of an alignment. Finally, we clean blocks within a MAF sequence alignment that reflect low-quality or paralogous alignments. We remove alignment blocks with length *<* 50 bases out of concern for alignment quality, and remove any remaining alignment blocks with paralogous structure (e.g. multiple sequences from the same species). Paralogous sequences refer to sets of sequences in the same species that are descended from the same ancestral sequence and were created as a result of gene duplication. Since our analysis focuses on Single Base Substitutions (SBS), we are not interested in the evolution of paralogous sequences and remove these alignment blocks from our final, cleaned MAF file (see Supp. Figure 9).

### 2.2 Inference of trinucleotide mutation rates

As is consistent with prior literature on somatic and germline mutational processes ^23,34–37^, we focus on the extraction of Single Base Substitutions (SBS) in the trinucleotide context to infer mutational processes. Inferring branch-specific mutation rates is a critical step in this procedure, and we do this on every terminal branch in our alignment (Figure 2b). For every species *s*, which has been aligned to reference genome *a_s_*based on evolutionary clade (see Figure 1), we split the alignment into 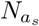 1Mb windows. For instance, four great ape species are aligned to the *Homo sapiens* reference genome, which we split into *N_Homo sapiens_* ≈ 2700 genomic regions. While previous studies on polymorphism data used 10Kb window sizes ^23^, we choose 1Mb windows to counteract the increased sparsity and noise of phylogenetic mutation data. Each row of our 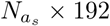 data matrix represents the observed mutation rate for the 192 trinucleotide mutation types in a given window (Figure 2c). Formally, we can define our data matrix for species *s* as 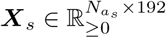 with 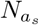 samples (genome windows) and 192 variables (trinucleotide mutation types) per sample.

**Figure 2:**
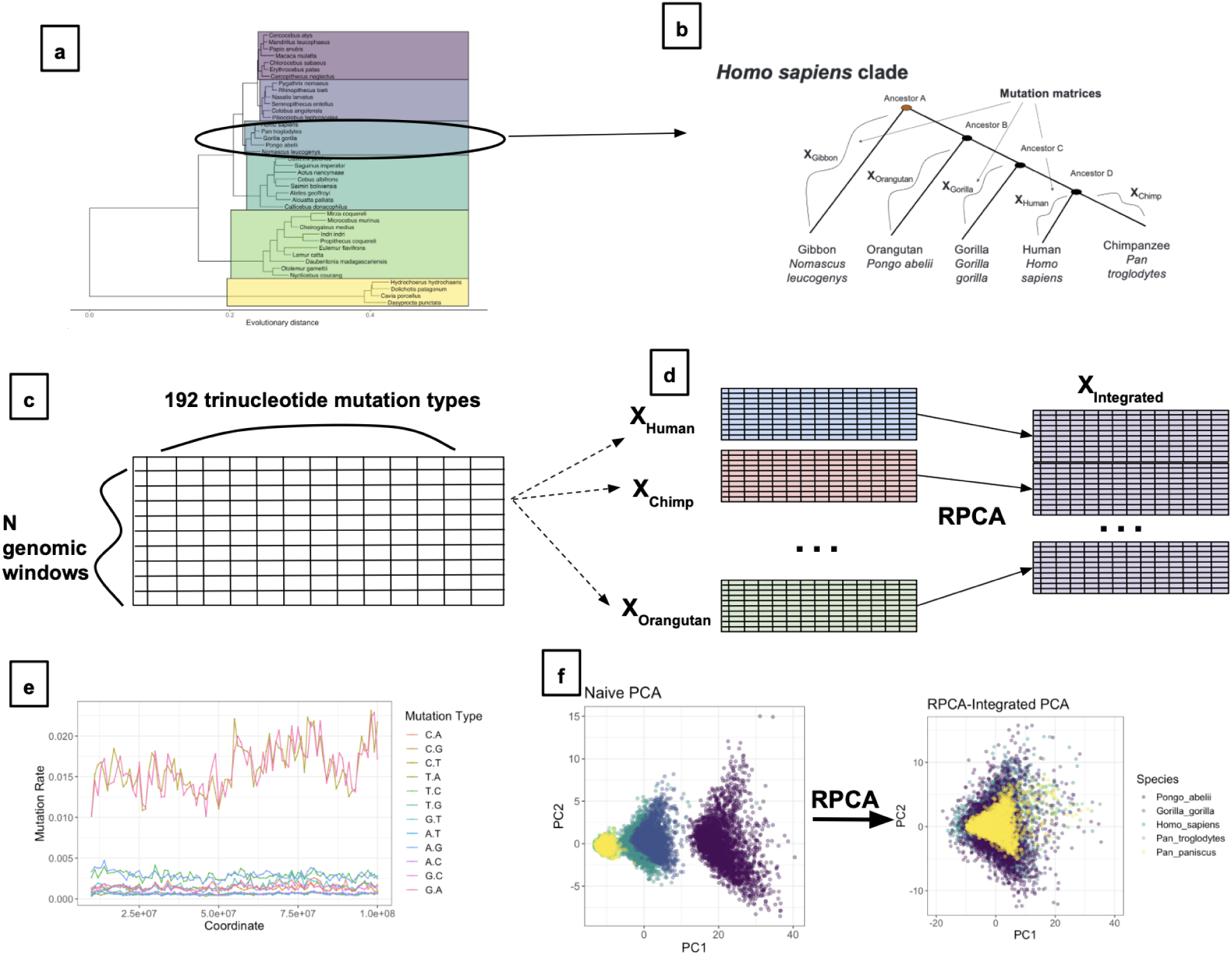
An overview of the data processing procedure. **a**.) Species are processed in groups based on reference sequence. **b**.) Within an evolutionary subclade (e.g. *Homo sapiens*), trinucleotide mutation rate matrices are inferred for every terminal branch. **c.**) Diagram of a species-specific trinucleotide rate matrices. One row contains trinucleotide mutation rates (192) for a 10MB region of the alignment. **d.**) Species-specific mutation rate matrices (*N_a_* × 192) are integrated via Reciprocal PCA into one, larger integrated rate matrix ((*N*_total species_*N_a_*) × 192). **e.**) Variations in single-nucleotide mutation rate can be witnessed across the genome. **f.)** PCA biplot of species within the *Homo sapiens* clade before RPCA integration (left panel) and after RPCA integration (right panel).

We then use nucleotide substitution models to estimate branch-specific mutation rates. Given our focus on extracting trinucleotide mutational processes, estimating these rates requires a substitution model with a trinucleotide context. Nucleotide substitution model selection remains an active area of research in evolutionary genetics ^38^, but these tools generally only consider single-nucleotide transition models ^39–41^. Trinucleotide models impose significant computational burdens because the interdependency of neighboring nucleotides means that computing the probability of transition for a single nucleotide requires consideration of all sites in a genomic window. Despite these obstacles, trinucleotide substitution models fit for shorter sequences ^42^, and versions that attempt to bound the propagation of nucleotide dependency ^43^. However, both of these implementations are still far too computationally intensive for our analysis, which requires estimating branch-specific mutation rates across 2 billion basepairs across multiple species.

In response to these shortcomings, we developed a novel approach to estimate branch-specific trinucleotide mutation rates that bypasses traditional ancestral state reconstruction. Our process splits the multispecies alignment into 16 context-specific subfiles, each representing sites with specific trinucleotide contexts. For example, the “A-N-G context subfile” contains all nucleotides (N) in the full alignment with an Adenine (A) adjacent to it in the 5’ direction and a Guanine (G) adjacent to it in the 3’ direction. Using PhyloFit ^42^, we estimate all 12 single-nucleotide substitution rates within each context subfile using the ‘unrestricted model’ (UNREST) ^44^. All possible single nucleotide substitutions have unique rates in an UNREST model. This accommodates asymmetric substution patterns characteristic of known germline mutational processes in humans ^23^. With all 16 context-subfiles, and their 12 respective single-nucleotide mutation rates, we can compute all 192 trinucleotide mutation rates. For example, we use the estimated C*>*T mutation rate in the the A-N-G subfile as a proxy for the ACG*>*ATG mutation rate for the whole alignment window. See Supplementary Appendix Section C for a more detailed overview of the mutation rate inference procedure.

Since context fixing removes overlapping dependencies, we avoid the computational burden of traditional context-aware nucleotide substitution models. Computing branch-specific trinucleotide mutation rates for 1 megabase windows across the entire *Homo sapiens* clade (Figure 2) can be parallelized and takes a few hours. We validate our procedure by comparing our estimated genome-wide mutational spectra on the *Homo sapiens* terminal branch with the mutation rates calculated from human polymorphism data ^23^ (Figure 13).

### 2.3 Signature extraction with Reciprocal PCA

We argue that the mutational process is the most appropriate biological level for studying the evolution of mutagenesis across species. Prior work has demonstrated that analyzing the variation in mutational spectra along the genome can effectively identify mutational processes from rare human polymorphisms ^23^. By extracting mutational processes–which we also refer to as “mutational signatures”–from mutational spectra across multiple species, we hope to elucidate both shared and species-specific mutagenesis patterns.

Previous research on mutational signatures has typically relied on Non-negative Matrix Factorization (NMF) for signature extraction ^34,35^, with more recent approaches employing volume-regularized variants of NMF with better convergence properties ^23^. By restricting mutational signatures and exposures to be non-negative, NMF imposes the assumption that observed mutational spectra are the result of additive combinations of *non negative* mutational processes. This assumption works well when studying somatic mutations in cancer because these mutations are often caused by distinct mutagens or cellular processes (e.g., exposure to UV radiation, tobacco carcinogens, or defective DNA repair) that produce specific mutation types in a straightforward, additive manner. However, enforcing this assumption on divergence data may lead to inaccurate conclusions ^45^. For example, the well-known process of Biased Gene Conversion (BGC) actively favors mutations of T or A bases over C or G bases ^46^. In this setting, a mutational signature model that allows for negative intensities would better reflect the promotion and suppression of different mutation types under BGC.

We find that even volume-regularized NMF performs quite poorly on our divergence-based substitution data. The increased noise in estimated trinucleotide substitution rates meant that NMF solutions were highly parameter dependent and inconsistent with known signatures from polymorphism-based studies ^23^. Instead, we use Reciprocal Principal Components Analysis (RPCA) and PCA as method of signature extraction. RPCA is commonly used to harmonize different experiments in single-cell RNA sequencing ^47^, and offers an identifiable, computationally tractable approach for jointly inferring signatures within an evolutionary clade.

Another significant weakness of a classical NMF-based approach is that all species must be modeled independently, even when the mutational processes in closely related species are likely quite similar. Early efforts to construct a “multi-study” NMF, which would allow for borrowing information between related species and could strengthen signature identification, are not yet computationally tractable for this study^48^. RPCA addresses this issue by integrating data from related species into one, anchored dataset. Since this anchored dataset is no longer non-negative, we then use Principal Component Analysis (PCA) to jointly extract signatures and exposures across an evolutionary clade ^36^.

RPCA leverages the fact that two related species likely have genomic regions with shared mutational spectra. Through identifying pairwise correspondences between pairs of genomic regions across species, called “anchors”, RPCA transforms species-specific data into a shared space. In this shared space, large-scale differences such as baseline mutation rate due to branch length have been accounted for (Figure 2f). Given data for two species, the RPCA integration procedure is as follows: 1.) project each species into the others’ PCA space, 2.) identify potential anchors through Mutual Nearest Neighbors (MNN)^49^, 3.) filter and weight these anchors based on the degree of similarity between the matched regions, and 4.) apply region-specific transformations, based on weighted combinations of anchors, to integrate the two datasets. To integrate multiple (*>* 2) species in the same clade, we follow the multiple dataset procedure outlined in Stuart et al. (2019), where the guide tree is constructed based on the observed similarity of species-specific datasets ^47^. In order to improve the quality of RPCA integrated datasets, we remove two species with low-quality mutation rate matrices (*Nomascus leucogenys* and *Dasyprocta punctata*), as determined by examining their raw PCA components.

At a basic level, an RPCA interpretation of dependence between a pair of species is binary. Two species’ respective mutation rate matrices can be combined or remain separate. We had already separated the original Zoonomia tree into subclades of closely related species to minimize problems associated with the projection of distant species onto the same reference genome (Figure 1). We use these same reference species clades to group species for RPCA integration. For example, all species aligned to the *Homo sapiens* reference genome (*Homo sapiens, Pan troglodytes, Gorilla gorilla, Pongo abelii*) are integrated together into 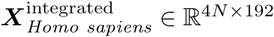. See Figure 2 for a high-level overview of the data processing pipeline and RPCA integration.

#### 2.3.1 Signature selection

After building the RPCA-integrated data matrix, we use PCA to extract the final signatures for each clade. Unlike NMF, PCA does not require choosing *a priori* a number of signatures *k* to extract. Instead, we filter the resulting principal components based on the “reflection test” ^23^. This test leverages the fact that DNA is a reverse-complementary molecule. A mutational process should either affect both DNA strands simultaneously, resulting in identical rates of reverse-complementary mutations, or operate on one strand at a time, leading to an anti-correlation of reverse-complementary mutations along the genome.

We calculate the reflection score for all inferred principal components, and plot this score as a function of principal component in Figure 3. We observe that earlier principal components tend to have higher reflection scores, followed by a steep dropoff in score between components 7-12. Given this observation, we choose to keep all signatures with reflection scores higher than 0.75. We present the number of components extracted per clade, and their average reflection scores in Table 1.

**Figure 3:**
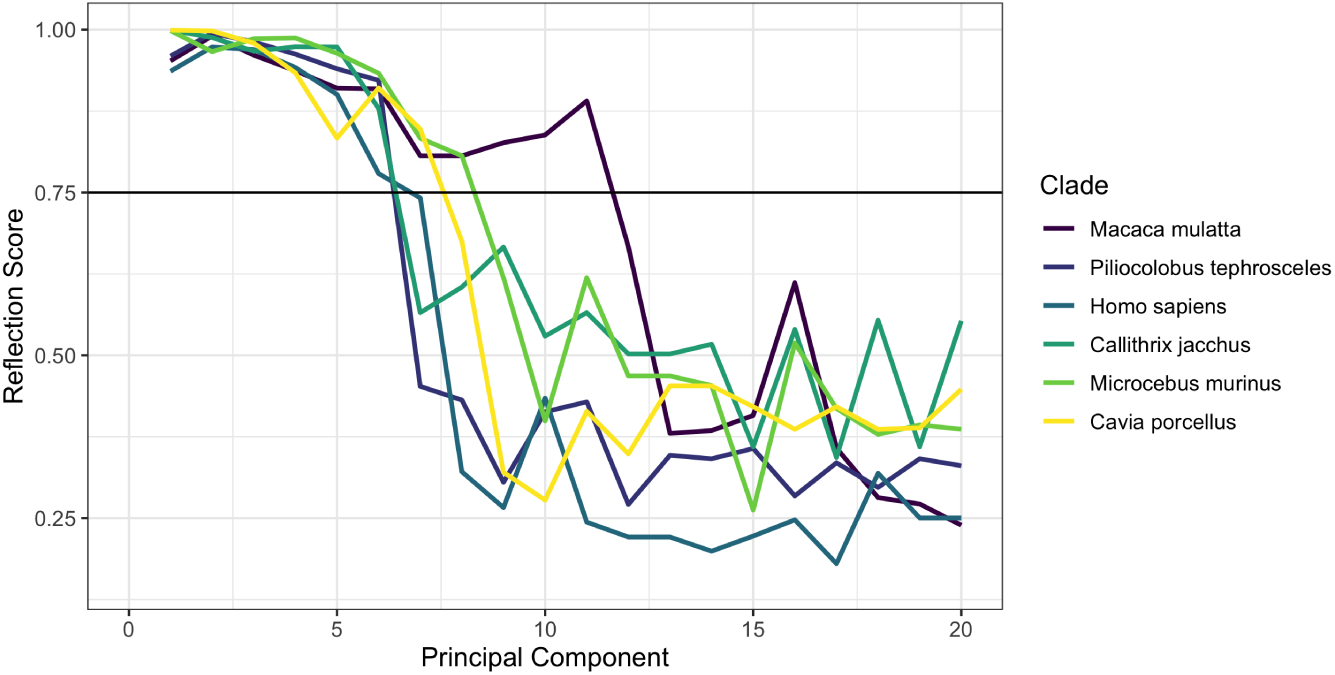
Component reflection score (y axis) vs component number (x axis) for all clades. The horizontal black line represents the reflection test cutoff.

**Table 1:**
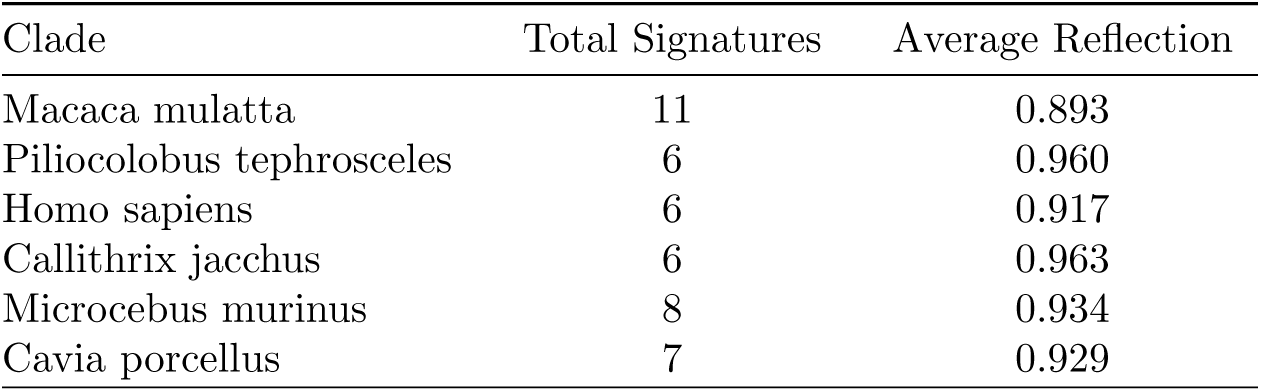
Number of signatures extracted per clade, and average reflection score among these signatures.

### 2.4 Validation with polymorphism data

Polymorphism-based mutational signatures represent an ideal case to validate our novel data processing and signature extraction procedure. The procedure to process divergence-based alignment data and polymorphism data has almost no overlap, so potential similarities in inferred signatures (and exposures) between the two approaches would be much more likely to represent real biological signal than a spurious artifact. To this end, we compare our divergence-inferred signatures and exposures with polymorphism-inferred signatures from humans, chimpanzees (*Pan troglodytes*) and baboons (*Papio anubis*). We use published signatures and exposures for our analysis of human polymorphism data ^23^. We process variant calls of chimpanzee and bonobo polymorphisms ^50^ and subsequently infer signatures with PCA (see Supp. Section D).

We find strong matches to *Homo sapiens* divergence processes 3, 4, and 6 in signatures inferred from human polymorphism data. These processes, extracted with orthogonal approaches on different datasets, have very similar profiles across the genome (Figure 4). We discuss the likely biological underpinnings of these processes in Section 3.

**Figure 4:**
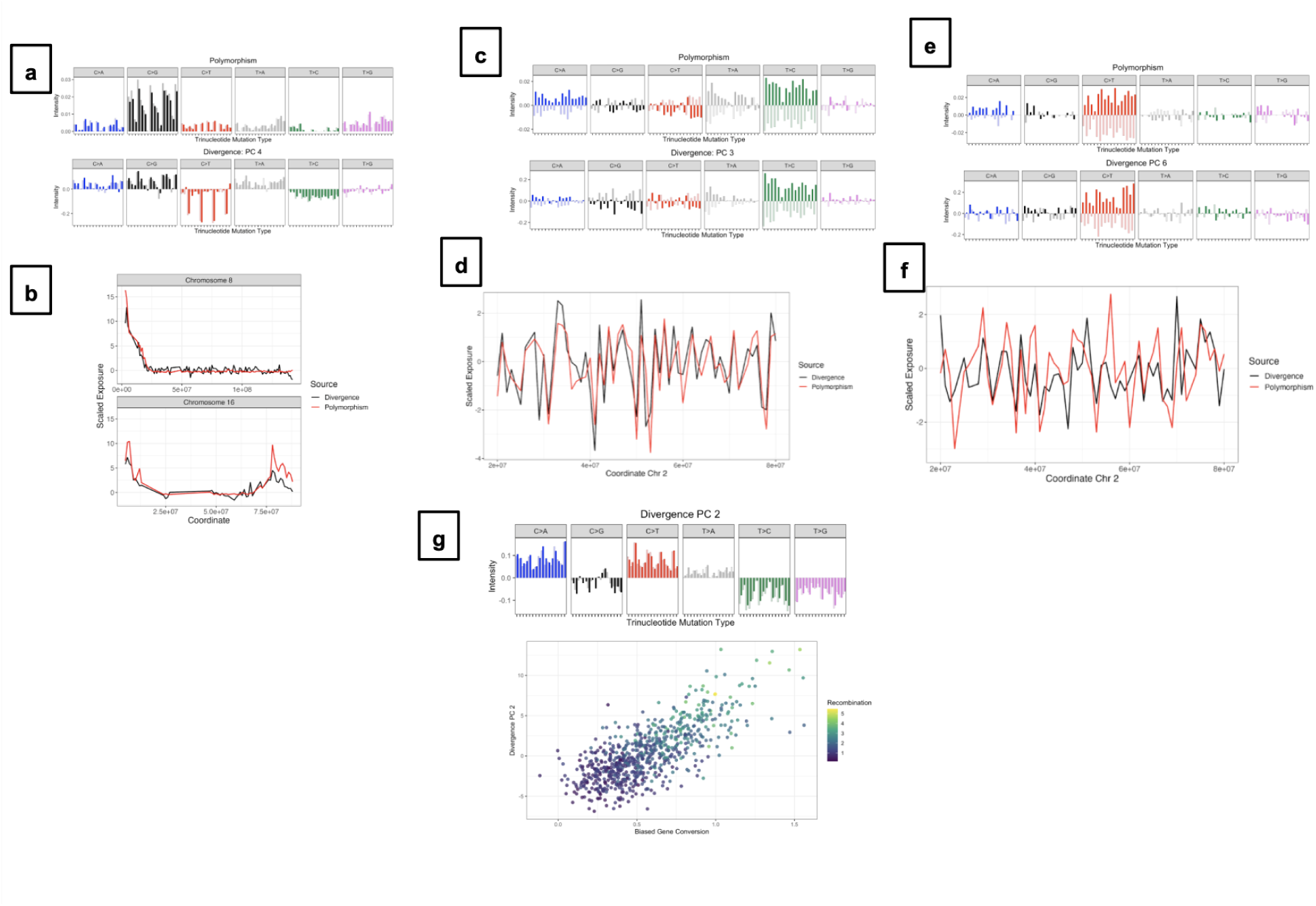
Comparison of polymorphism- and divergence-based signatures and exposures. In figures **a**, **c**, and **e**, polymorphism-inferred signatures are in the top panel and divergence-inferred signatures are in the bottom panel. In figures **b**, **d**, **f**, red lines represent exposures from polymorphism processes and black lines represent exposures from divergence processes. **a.)** “Maternal” C*>*G signature, with exposures over **b.)** hotspots on Chromosome 8 and Chromosome 16. **c.)** Bulky lesions (asymmetric errors with transcription), and **d.)** exposures over coordinates on Chromosome 2. **e.)** C*>*T signature with **f.)** correlation in exposure over coordinates on Chromosome 2. **g.)** Trinucleotide spectra of the BGC signature (top panel). Bottom panel, BGC process exposure (y-axis) vs known BGC activity (x-axis) across the human genome. Color gradient represents the known recombination rate, with brighter colors representing higher recombination rates.

We also validate our data processing and RPCA-based signature extraction with polymorphism data from chimpanzees (*Pan troglodytes*) and baboons (*Papio anubis*). Due to the difference in reference sequence alignments, we focus on the similarity between inferred signatures instead of exposures. All *Homo sapiens* clade divergence signatures have a strong match with polymorphism-based signatures (Figure 5a). Two of these matches, which we refer to as Biased Gene Conversion (Figure 5b) and bulky lesions (Figure 5c), are shown in the main text (see Supp. Figure 14 and 15 for full sets of signatures). Similarly, we also find a high degree of similarity between *Macaca mulatta* clade RPCA signatures and *Papio anubis* polymorphism signatures (Figure 5d). The *Macaca mulatta* clade contains twice the number of species as the *Homo sapiens* clade, and we were accordingly able to extract significantly more macaque-clade signatures (11 vs 6). However, due to sample size and data quality limitations with *Papio anubis* polymorphism data, we were only able to extract eight high-quality polymorphism signatures. While this makes it difficult to validate all divergence-inferred signatures like in *Pan troglodtytes*, we still find that the biased gene conversion and bulky lesions signatures are highly similar between the two data sources.

**Figure 5:**
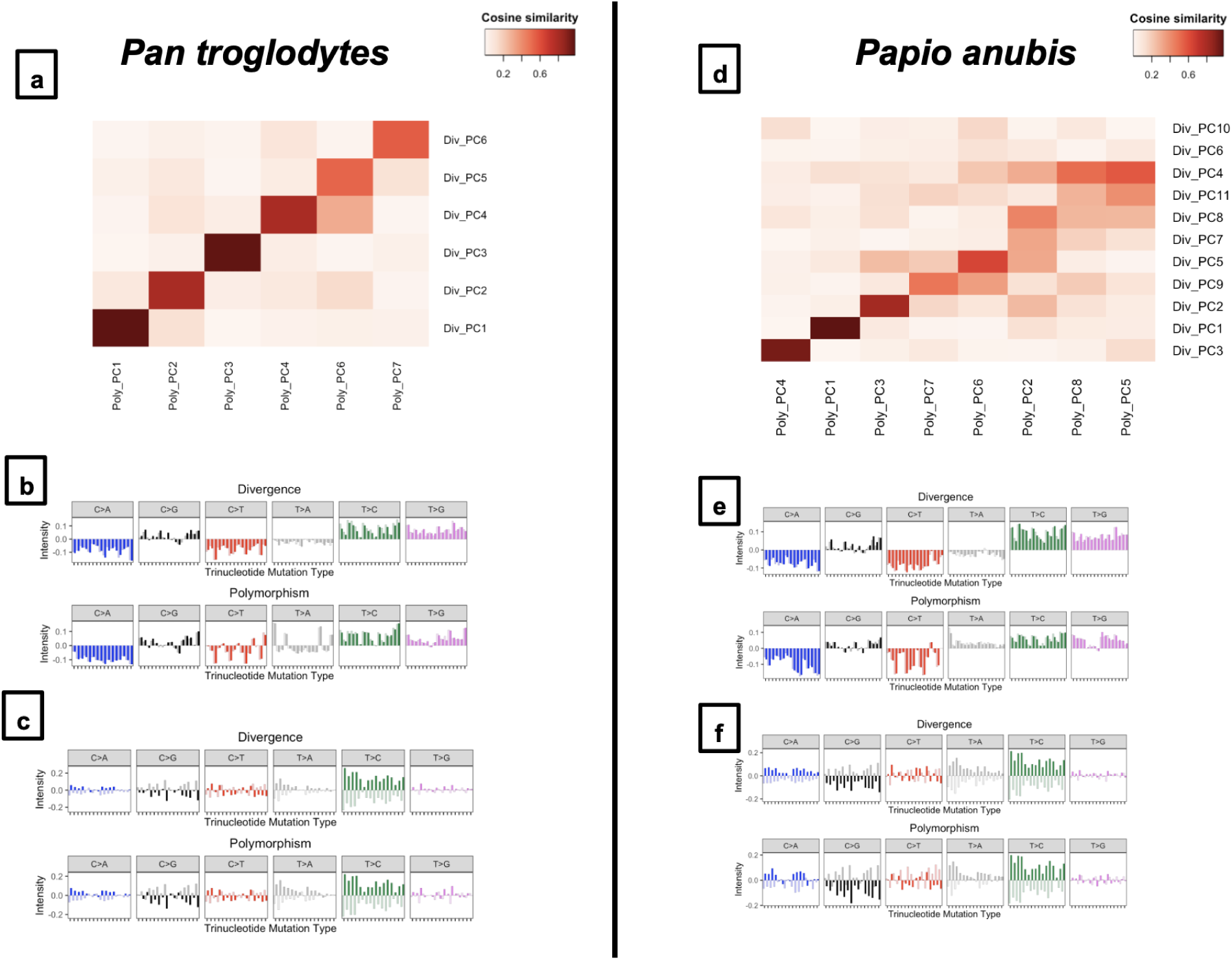
Comparison of signatures inferred from divergence- and polymorphism-based data. **Left panel:** Comparisons with *Pan troglodytes*. **Right panel:** Comparisons with *Papio anubis*. **a.)** and **d.)** Heatmap of correlations between polymorphism-inferred signatures (columns) and divergence-RPCA inferred signatures (rows). **b.)** and **e.)** Biased gene conversion signature, extracted from divergence (top panel) and polymorphism (bottom panel). **c.)** and **f.)** Bulky lesions signature, from divergence (top panel) and polymorphism (bottom panel).

## 3 Results

We now overview the mutational signatures found across our studied primate species. After implementing our signature selection criteria (section 2.3.1), we extract 44 total signatures that pass the reflection test across the six studied evolutionary clades. The breakdown of signatures per clade is summarized in Table 1. We then concatenate this collection of signatures into a matrix ***S*** ∈ ℝ^44^*^×^*^192^, and compute the signature-signature similarity matrix 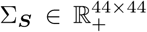 based on cosine similarity, a common method of signature comparison ^45^. In this notation, [Σ***_S_***]*_ij_* = |Cos(***S****_i_,* ***S****_j_*)|. Figure 6 provides a visualization of this signature-signature correlation matrix. The clusters of similar signatures on Figure 6, characterized by dark red squares, suggest widespread conservation of certain germline mutational processes across primates.

**Figure 6:**
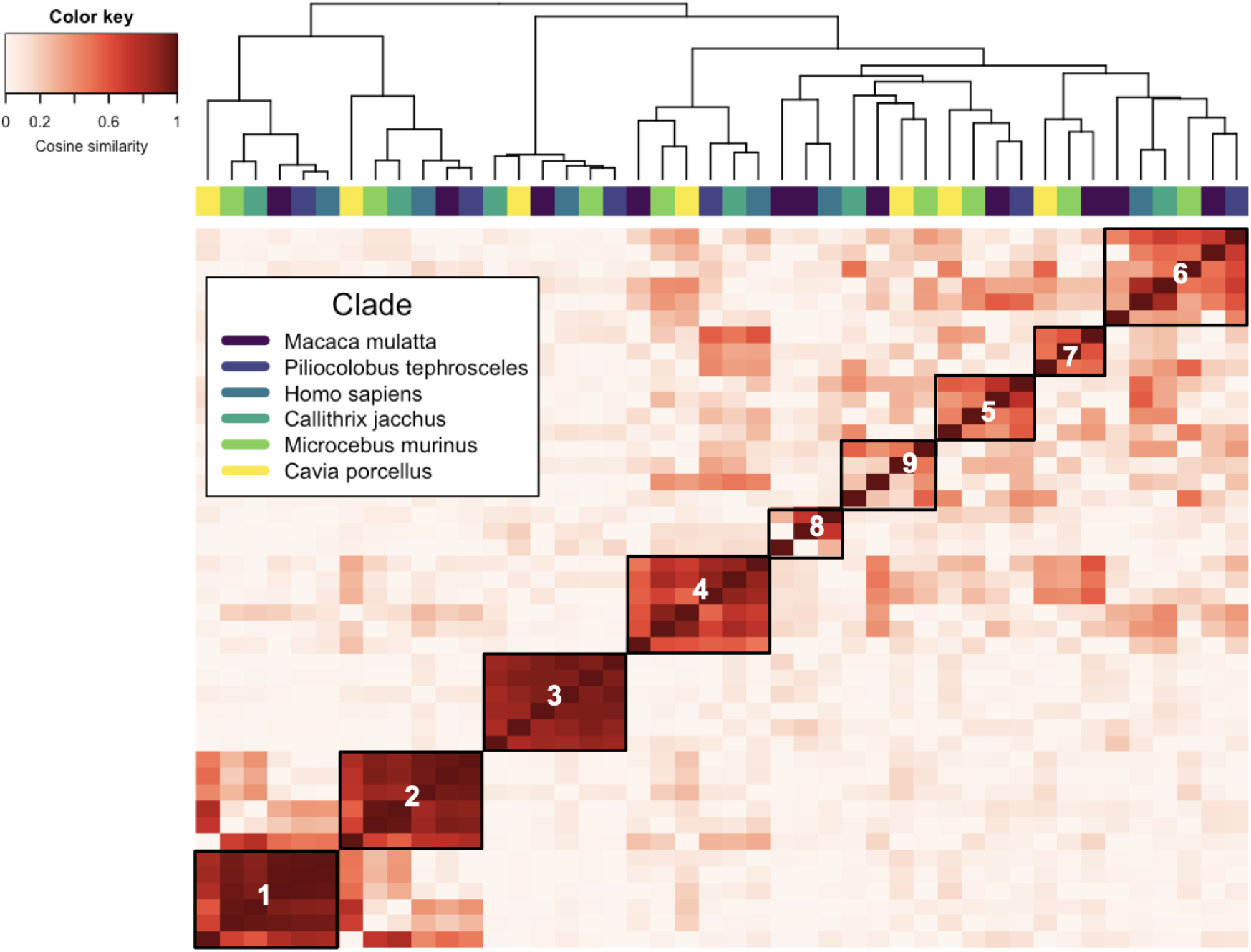
Heatmap of the signature-signature correlation matrix. Color intensity at row *i* and column *j* represents the cosine similarity between process *i* and process *j*. Colors above the heatmap indicate the clade that the signature belongs to. The dendrogram indicates the high-level clustering structure of the signatures. Numbers written on the heatmap refer to the cluster assigment, i.e. the signatures within the box labeled “1” belong to cluster 1 from Figure 7.

We perform hierarchical clustering with complete linkage on the signature-signature correlation matrix to find clusters of similar signatures. Choosing 9 clusters both allows for a minimal within-cluster sum of squares and a high within-cluster silhouette score (Supp. Figure 16). We compute the centroids for each of these clusters and display them in Figure 7. Table 2 describes the reflection scores of the centroids and average silhouette scores per cluster.

**Figure 7:**
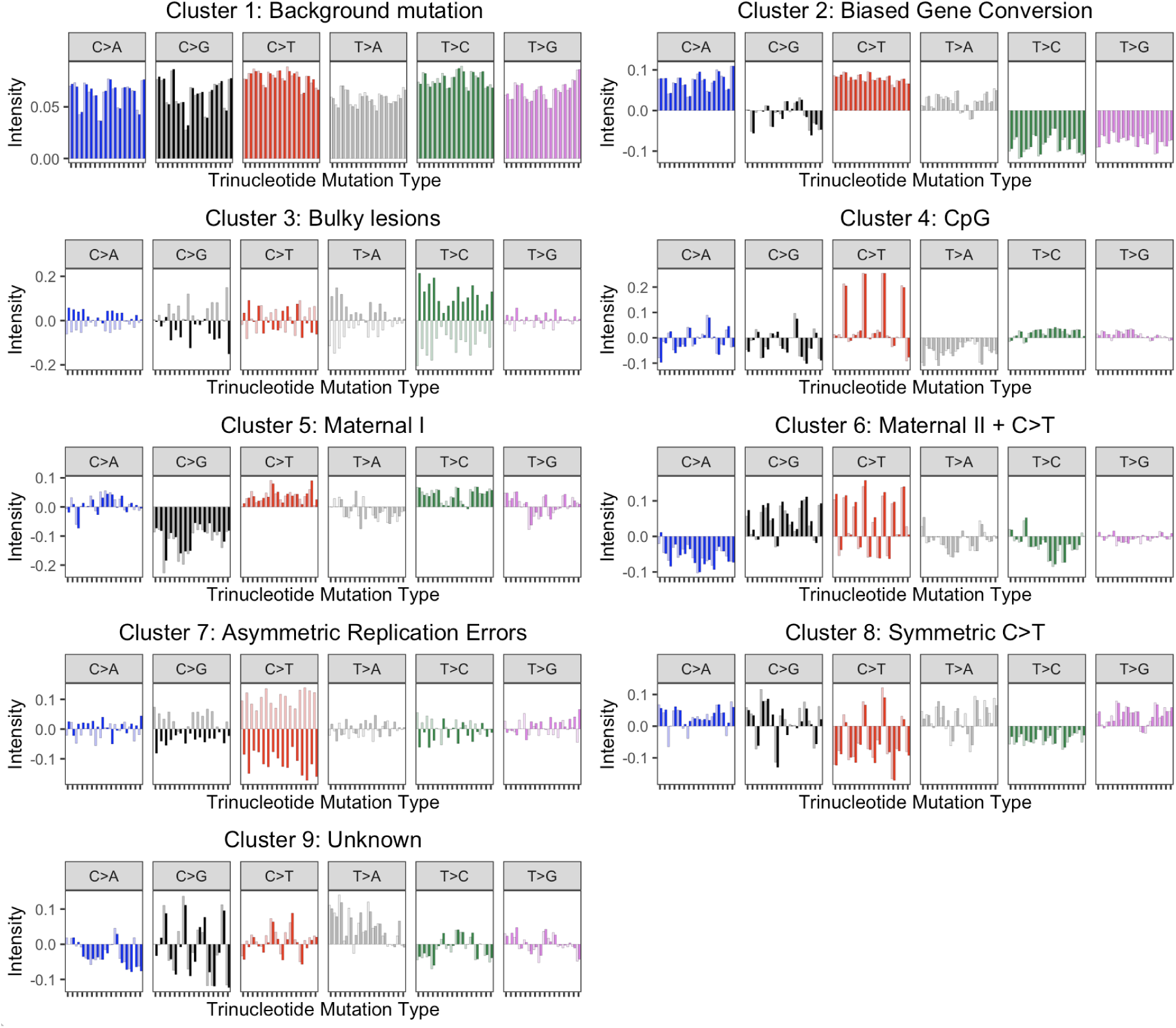
Centroids of all nine mutational process clusters.

**Table 2:** Selected statistics on the nine inferred mutational process clusters.

| Cluster | Cluster Name | N. Processes | Reflection | Silhouette |
| --- | --- | --- | --- | --- |
| 1 | Background mutation | 6 | 0.984 | 0.636 |
| 2 | Biased Gene Conversion | 6 | 0.998 | 0.603 |
| 3 | Bulky lesions | 6 | 0.994 | 0.850 |
| 4 | CpG | 6 | 0.990 | 0.423 |
| 5 | Maternal I | 3 | 0.954 | 0.375 |
| 6 | Maternal II + C>T | 6 | 0.979 | 0.248 |
| 7 | Asymmetric Replication Errors | 3 | 0.921 | 0.313 |
| 8 | Symmetric C>T | 4 | 0.939 | 0.339 |
| 9 | Unknown | 4 | 0.929 | 0.125 |

Processes in cluster 1 represent background mutation. This process is observed in all evolutionary clades, and is characterized by roughly equal intensities from all trinucleotide mutation types. Thus, a change in the exposure to a cluster 1 process would be equivalent to change in the overall mutation rate for a given genomic region.

Processes in cluster 2 are characterized by increases in the rate of T*>*C and T*>*G mutations and decreases in the rate of C*>*G and T*>*A mutations. This mirrors the process of “Biased Gene Conversion” (BGC), where DNA repair after recombination tends to prefer the more thermodynamically stable Guanine and Cytosine nucleotides over Adenine and Thymine ^46^. Figure 4 demonstrates a strong correlation of the exposure to this process with published metrics of BGC strength (Cor = 0.73) and recombination rate in humans ^51^.

Cluster 3 represents a closely related group of “asymmetric” mutational processes which are present across all six evolutionary clades. When a genomic region has a positive exposure to this process, there is an increase in T*>*C mutations. On the other hand, a negative exposure corresponds to an increase in complementary A*>*G mutations. This directly corresponds to the action of transcription-coupled repair, which depletes A*>*G mutations on the transcribed strand ^52^. The intensity of a similar process from polymorphism-based data has been shown to correlate with the direction of transcription ^23^, and we notice tightly correlated exposure patterns between the divergence- and polymorphism based processes along coordinates in Chromosome 2.

Processes in cluster 4 are characterized by CpG transitions. Like clusters 1-3, this cluster contains processes from all six evolutionary clades. We do not find a distinct CpG transversions process from our procedure, although some processes (cluster 6 and cluster 9) demonstrate some promotion of CpG transversions.

Processes from clusters 5 and 6 demonstrate strong similarity to known processes of “maternal mutagenesis” in humans. This signature was previously shown to co-localize with regions prone to mutation accumulation in the maternal germline ^23,53^. The divergence-inferred signatures demonstrate similar patterns of C*>*G transversions to the polymorphism-inferred signature (Figure 4a), and we compare their activities in known hotspot regions on Chromosome 8 and 16 (Figure 4b).

The asymmetric processes in cluster 7 are characterized by C*>*T mutations. Exposures to this process in *Homo sapiens* correlate well with polymorphism-inferred exposures to a similar process that has been connected to errors stemming from DNA replication ^23^. Processes in cluster 8 are also dominated by C*>*T mutations, but do not exhibit any asymmetry. We hypothesize that processes in these two clusters are related, a product of our data processing pipeline where we calculate trinucleotide mutation rates over relatively large (1Mb) windows compared to the 10Kb windows used polymorphism-based studies ^23^.

Cluster 9 processes are characterized by T*>*A mutations, as well as spikes in CpG transversions. This cluster does not match any previously detected germline processes, has the lowest silhouette score out of all clusters, and could not be validated in any of our three polymorphism-based datasets. These processes may represent a genuine biological process, or may be the result of an artifact from our data processing procedure.

In addition to comparing inferred processes across evolutionary clades, we also study the evolution of exposures to these processes within a clade. Comparisons of exposures are made more feasible because we extract trinucleotide mutation rates against a common reference sequence within a clade, and use RPCA to resolve systemic differences between species (e.g. variations in overall mutation rate due to branch length). We then note that with PCA, a component’s “importance” for explaining variation in the overall data is directly proportional to the variance of its principal component scores. Using this interpretation, we use the species-specific observed standard deviation of exposures (i.e. a vector of PC scores) for a certain process (a PC component) as our metric of process “strength” within a species. We also refer to this as the process “intensity” for a given species.

Figure 8 displays our comparison of genome-wide process importance between species. With the exception of components 4-9 in the *Macaca mulatta* clade, process strength appears remarkably consistent across species. A few species (e.g. *Pongo abelii: Homo sapiens* clade, *Nycticebus coucang: Microcebus murinus* clade) have larger process intensities across all components. This suggests a residual difference in overall mutation rate that RPCA was unable to resolve, rather than a process-specific switch within a species. Since these species also have relatively long terminal branch lengths, it is likely that these differences are an artifact of our data processing procedure and RPCA integration rather than a real biological signal.

**Figure 8:**
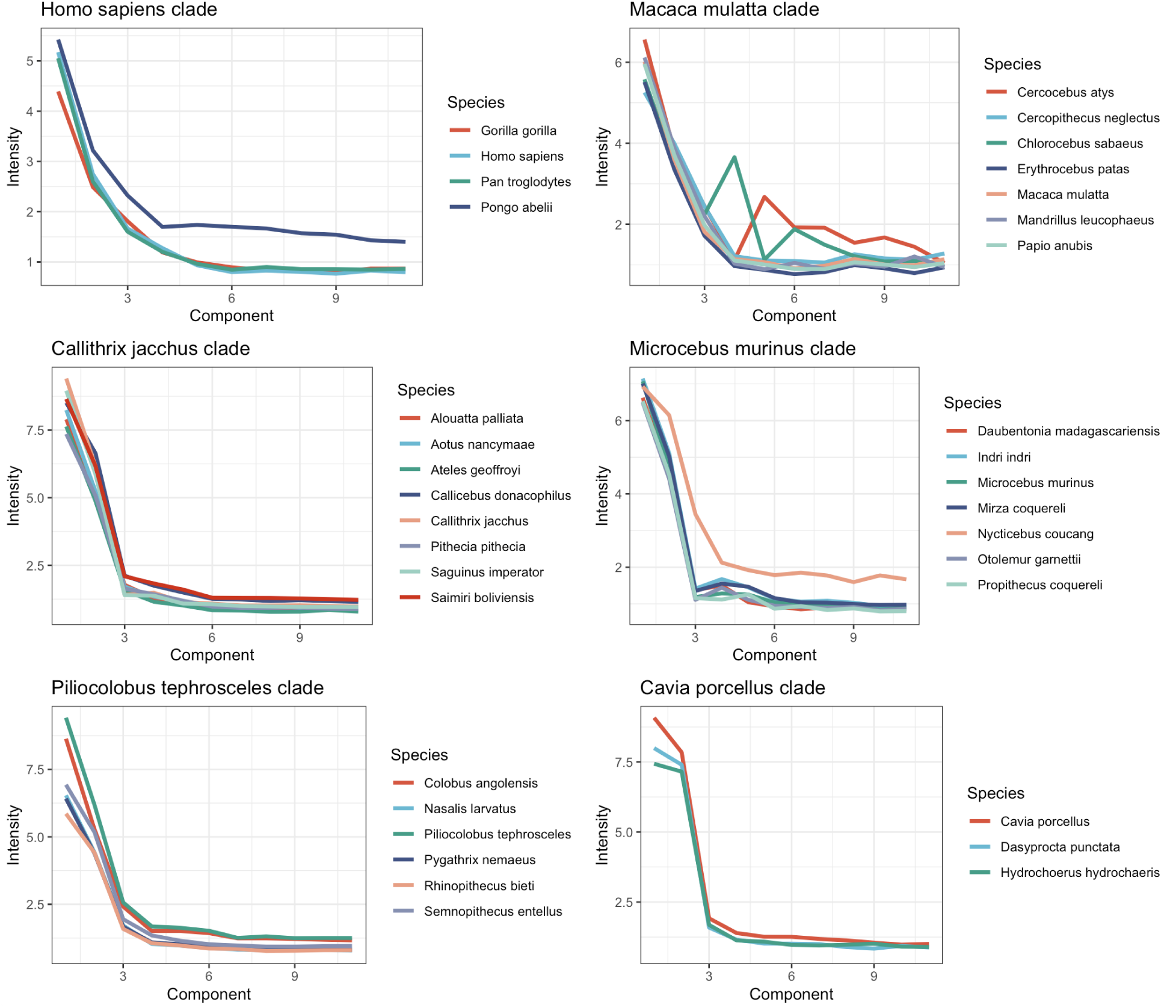
Standard deviation of principal component score, by principal component, for individual RPCA-integrated clades.

## 4 Discussion

This work provides a novel, spatial investigation of mutational processes beyond the single-nucleotide context in non-human species. We extract fourty-four mutational processes across six independent phylogenetic clades, which we group into nine clusters of processes. Of these nine clusters, we suggest plausible biological explanations for eight of them. While the mechanisms behind these mutational processes are not fully understood, their prevalence throughout phylogenetic history makes a strong case for their validity as “true” biological signatures. These signatures also appear in independently processed polymorphism data and correspond with previously published exposures in humans ^23^, providing further evidence for their biological validity.

We find strong evidence for the presence of mutational processes related to background mutation, biased gene conversion, asymmetric repair of bulky lesions, and CpG mutations across all six evolutionary clades. Since our selected clades include five primate clades, as well as a small outgroup of hamster-related species (*Cavia porcellus* clade), it is likely that these processes persist across a large subset of mammal species.

We also find processes connected to maternal mutagenesis in all primate clades but not the hamster (*Cavia porcellus*) clade. While the hamster clade does contain a process characterized by an increase in C*>*G mutation types, this process appears visually different than those found in primate clades as it is less dominated by CpG transversions. Since cluster 5 agrees with evolutionary relationships (e.g. all primates have the maternal process, while hamsters do not), it is possible that the maternal process lines up with a biological life trait that has been lost in hamsters. However, it is possible that the maternal process is ubiquitous across studied clades but is subject to detectability issues with PCA. For example, we extract processes related to asymmetric replication errors in both *Macaca mulatta* and *Homo sapiens* clades, but not the *Piliocolobus tephrosceles* clade which is sister to *Macaca mulatta*. While it is possible that the process was “lost” in the *Piliocolobus tephrosceles* clade, it is more likely that our signature extraction procedure was unable to detect the process because of noise or errors in the original alignment.

We also identify the trinucleotide signature of Biased Gene Conversion. Prior research, which used the spectra of rare Single Nucleotide Polymorphisms (SNPs) to infer mutational signatures, did not identify a biased gene conversion signature ^23^. Rare SNPs have not yet been exposed to many rounds of recombination and the cumulative effects of BGC, whereas the common variants found in divergence data would have been highly exposed to BGC processes over millions of years of evolution. The identification of a trinucleotide BGC signature opens the door for future investigations into the process’ evolution across mammals.

In order to accommodate the unique properties of phylogenetic sequence data, we develop an innovative and efficient approach to infer branch-specific trinucleotide mutation rates while considering all possible ancestral states. Existing method for mutation rate estimation beyond the single nucleotide level are incredibly computationally intensive due to the propagation of statistical dependency along the genome ^4,42,43^. Our key insights come from restricting the propagation of dependency in nucleotide transitions to the immediate 5’ and 3’ nucleotides, as well as inferring the identity of neighboring ancestral states with parsimony when possible. We compare our estimated trinucleotide mutation rates on the *Homo sapiens* terminal branch with polymorphism-inferred mutation rates, and find large similarities. While this methodology is not entirely bias-free, it allows us to build the species-specific mutation rate matrices at a reasonable computational scale.

### 4.1 Limitations

One of the most challenging aspects of using phylogenetic sequence data for mutational signature extraction is the sparsity of mutations. Compared to the mutation rate matrix obtained from human polymorphism data ^23^, our mutation rate matrix inferred from Cactus data on the *Homo sapiens* terminal branch has an order of magnitude increase in sparsity. In order to counteract the increase in sparsity and other phylogenetic noise factors that are prevalent at smaller smaller genomic scales, we use 1MB window sizes for this analysis. While this allows us to achieve cleaner solutions compared to smaller window sizes, it also prevents us from extracting mutational processes that are only variable at smaller scales and are undetectable in 1MB windows. For example, a strand-dependent maternal mutational process is detectable at the 10KB scale in polymorphism data but we can only extract a strand-dependent maternal process from 1MB Cactus data. Additionally, smaller genomic windows could have provided more insightful comparisons of signatures for similar reasons. CpG islands, which have been shown to have higher rates of CpG transversions and lower rates of CpG*>*TpG mutations, predominantly exist below the 1MB scale and so this relationship between signatures is less detectable in our analysis.

While our signature extraction procedure with RPCA integration and PCA allows us to extract stable and biologically interpretable signatures, it makes no guarantees about signature detectability. Knowing that a mutational process is present in a clade does not guarantee that PCA will detect it. We notice this with the extraction of processes related to replication timing, hypothesizing that the lack of such a process in the *Piliocolobus tephrosceles* clade is due to detectability issues rather than a real biological phenomenon. Extracting species jointly within a phylogenetic clade carries the assumption that these species have shared or closely related mutational processes, which may not always be true. The results of this study suggest this assumption may be reasonable, as we find widespread conservation of processes across clades where signature extraction happened independently. However, a more flexible model structure that explicitly allows for species-specific signatures ^48^ could be advantageous for future work.

### 4.2 Future directions

This work uncovers the biology of mutagenesis across more than 90 million years of evolution and offers a framework for future investigations of mutational processes development. Limited data exists on DNA features of non-human species ^54^, and signatures extracted from this approach can be used as a proxy for features such as the direction of replication, replication timing, or repair of bulky lesions. Furthermore, the evolution of mutational processes can be studied based on changes in process exposures along the genome or on changes in the process spectra itself. By using process intensity as a proxy for phenotype on the tree, genetic changes underlying this process evolution could be identified using modern phylogenetic “gene to phenotype” methods ^55,56^. For example, the evolution of Biased Gene Conversion, which we identify with a trinucleotide signature in this work, can be studied with these PhyloG2P methods. Other factors such as environment, generation time or branch length could be analyzed for their impact on mutagenesis. We also develop a novel, computationally efficient trinucleotide mutation rate estimation model that can be used for other context-aware phylogenetic inference. We demonstrate a robust procedure for inferring germline mutational processes across 40 mammalian species which can easily be expanded to other species with high-quality alignments.

## A Supplementary Appendix

## B MAF file cleaning procedure

**Figure 9:**
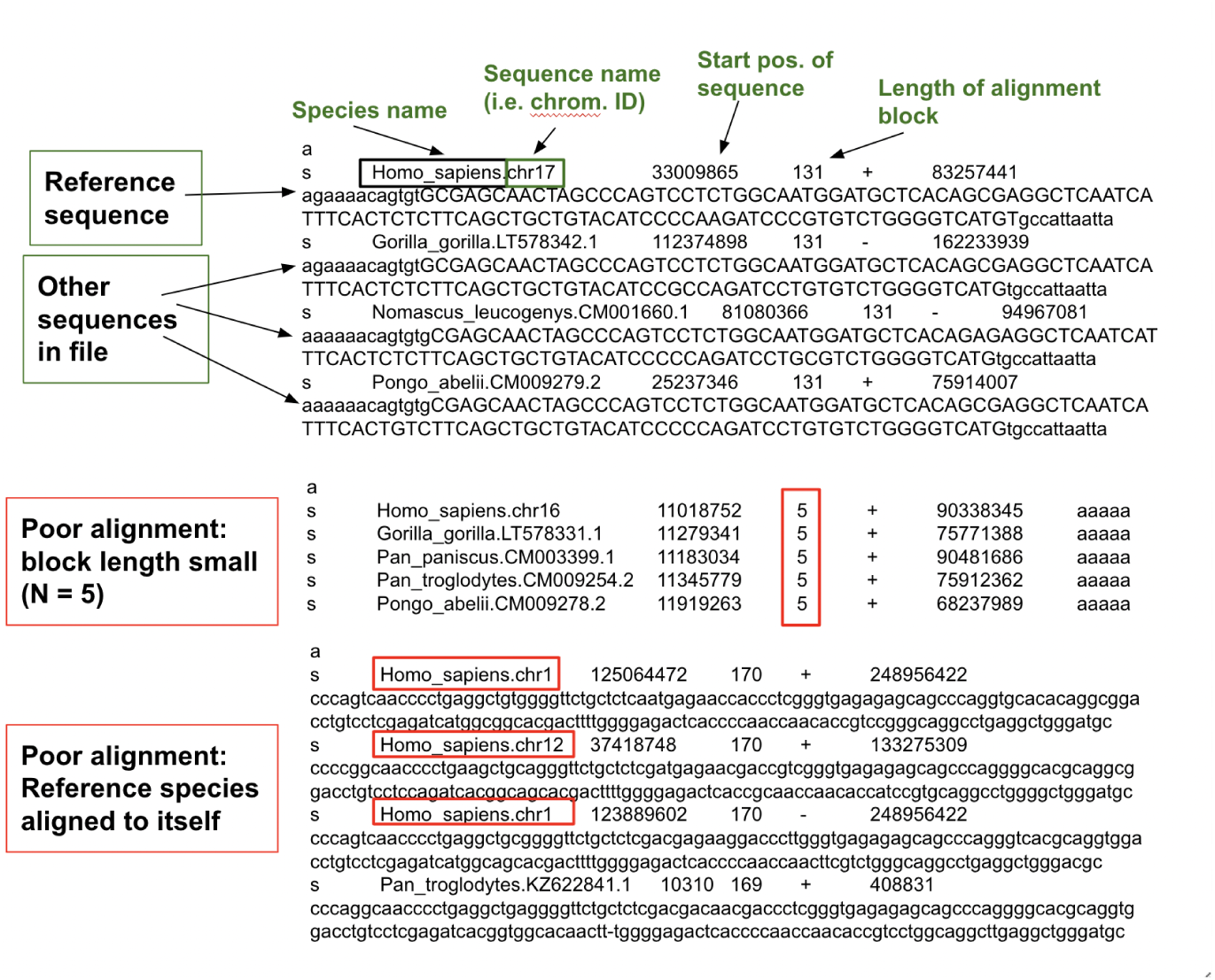
A diagram of the MAF file format with three separate alignment “blocks”. The first alignment block represents a clean sequence alignment to *Homo sapiens* chromosome 17. We remove the last two blocks from the final MAF file because of a low total block length or because of paralogous sequence.

## C Inference of Trinucleotide Mutation Rates

Inferring branch-specific mutation rates is a critical step in our procedure. Starting with aligned, cleaned MAF files, we build a mutation rate matrix for every species in the alignment. We split the aligned genome *a* into windows of length *b*, creating *N_ab_* total alignment windows. In order to counteract the increased sparsity and noise of phylogenetic mutation data, we choose window length *b* = 10^6^. Each row of our *N_ab_* × 192 data matrix represents the observed mutation rate for the 192 trinucleotide mutation types in a given window. Formally, we can define our data matrix for species *s* as 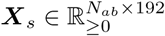 with *N_ab_* samples (genome windows) and 192 variables (trinucleotide mutation types) per sample. This can be written in matrix form as:

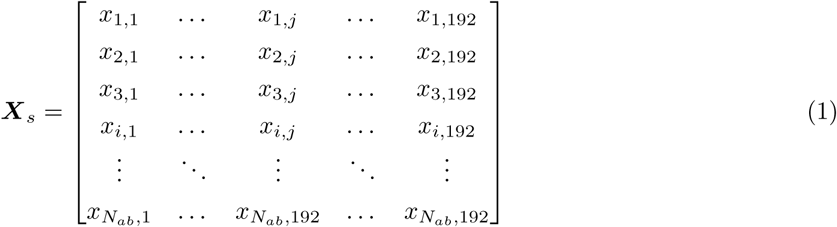

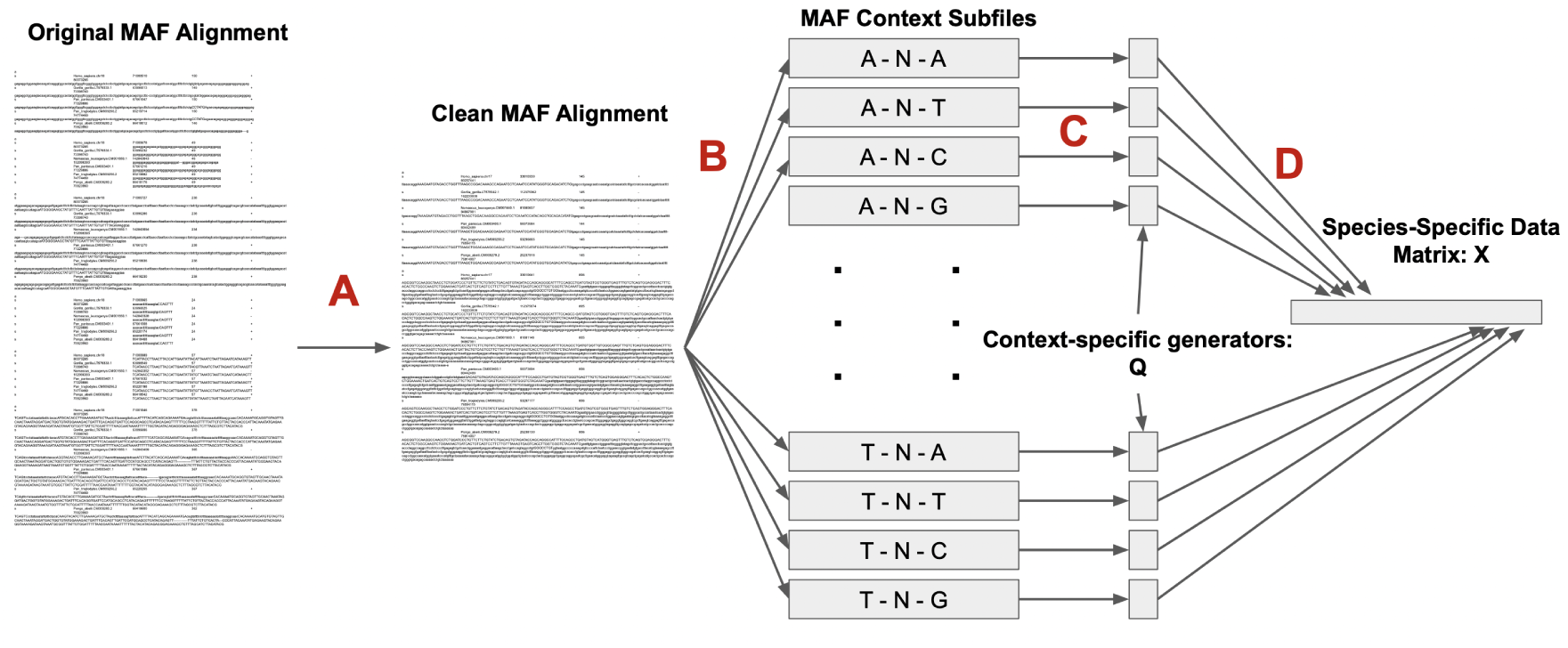
(a) The full MAF to species-specific data matrix ***X****^s^* processing pipeline. **A**: Clean short alignment blocks and paralogous alignments from original MAF file. **B**: Split cleaned MAF alignment into 16 context-subfiles for every alignment window. **C**: Using PhyloFit, infer the species-specific generator matrix 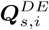. **D**: Scale every element of 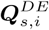, by the estimated branch length 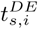. If necessary, average generator matrices across PhyloFit runs. Combine the 16 generator matrices, with 12 parameter values each, into one row of ***X****^s^*.

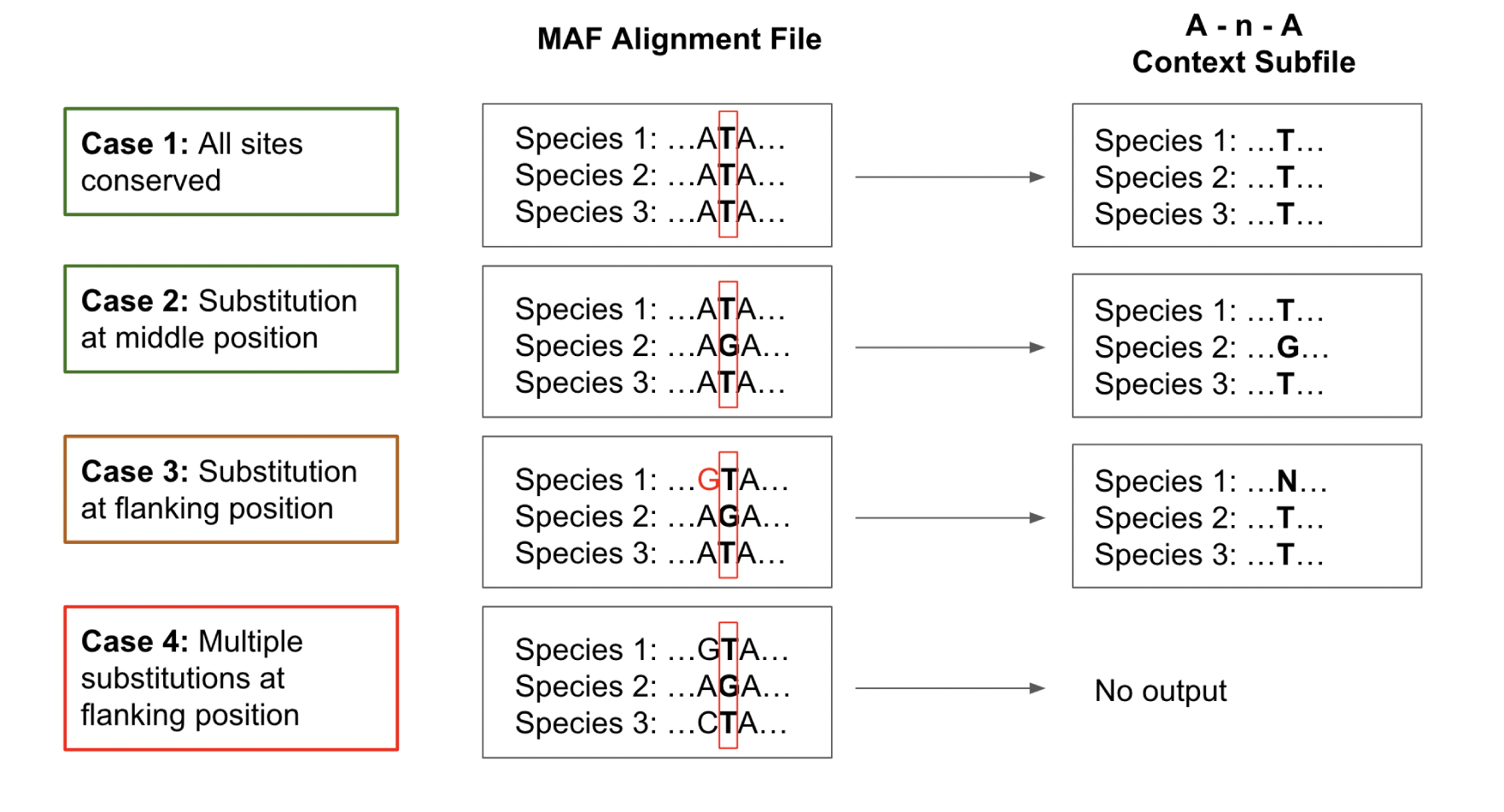
(b) Diagram of the rules for the creation of the MAF context subfiles.

We arrive at this data matrix ***X****_s_* for every terminal branch in a given alignment (see Figure 11).

### C.0.1 Problem: Latent Ancestral States

In order to build the species-specific mutation matrix, we need to assign mutations to different phylogenetic branches. In order to do this, we must model the latent, ancestral states. Unlike the use of polymorphism data in **(author?)** ^23^, which relies on a human reference genome to determine the ancestral and corresponding derived states at every basepair, identifying ancestral states with phylogenetic data is much more complicated and is an area of active research ^57,58^. With the Cactus alignment we observe the nucleotide sequence of all terminal nodes in the data. However, since we do not know the exact genomic sequence of the ancestral nodes, we must use statistical models to estimate them (see Figure 12). As most previous research in evolutionary genetics has focused on single-nucleotide mutation rates, our consideration of trinucleotide mutation rates requires the development of a novel data-processing pipeline.

**Figure 11:**
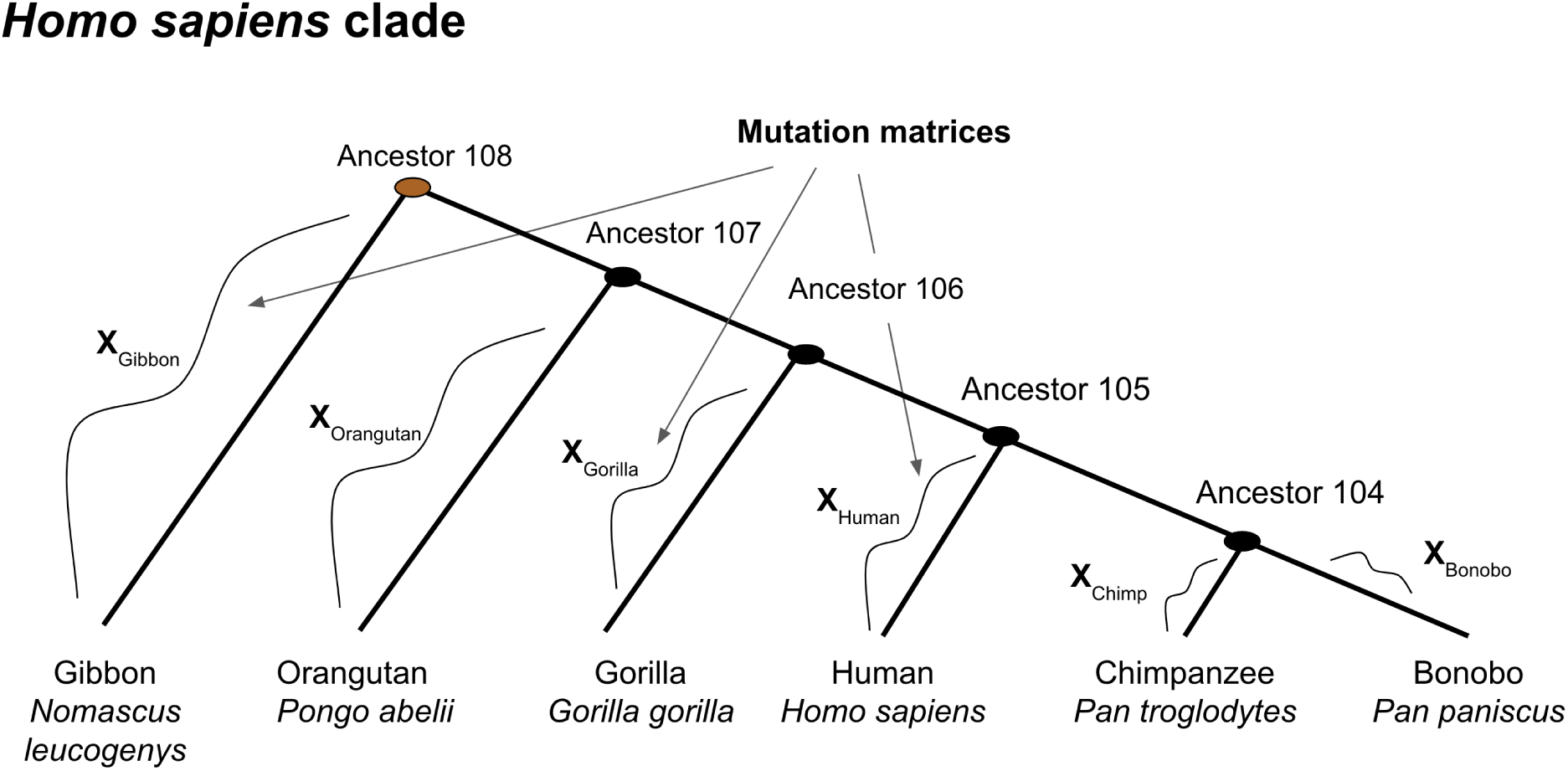
The mutation rate matrices inferred based on the phylogenetic tree for the “human” clade. We infer separate mutation rate matrices for all terminal branches.

**Figure 12:**
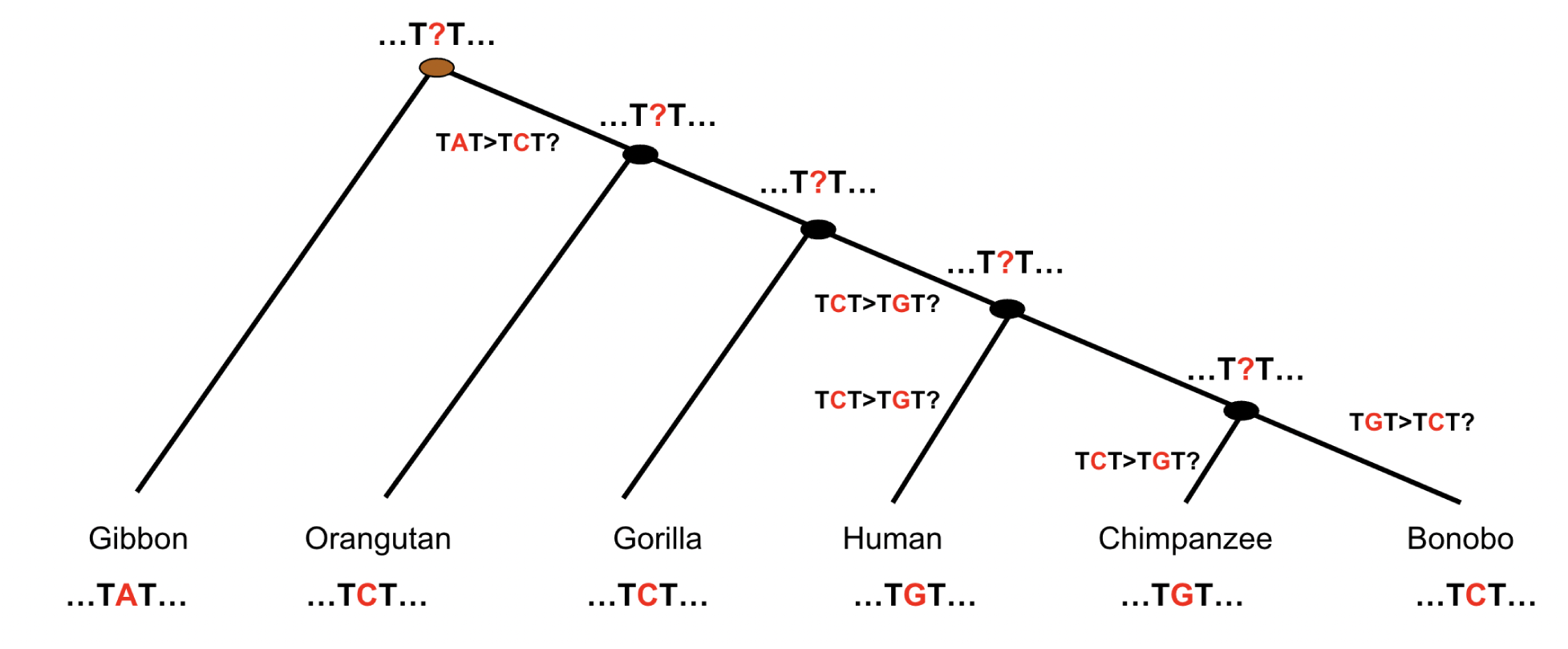
For the pattern of 3-mers observed in the terminal nodes of the *Homo sapiens* clade, we must incorporate estimations of the ancestral states in order to place mutations on phylogenetic branches.

The Cactus alignment provides estimated ancestral sequences, but the model used to produce these sequences makes the strong simplifying assumption of uniform nucleotide transition rates. The Cactus ancestral state reconstruction procedure uses a Jukes-Cantor model ^59^. The simplest nucleotide substitution model, the Jukes-Cantor model assumes that all base pairs mutate at the same rate. By enforcing that all nucleotide transitions are equally likely, the Jukes-Cantor model ignores well-established differences in nucleotide-specific mutation rates. CpG sites, where a Cytosine nucleotide is followed by a Guanine nucleotide in the linear sequence of bases, mutate to TpG at an order of magnitude higher rate than any other pair of nucleotides. Because the Jukes-Cantor model does not account for the increased rate of CpG mutations, it is underspecified and will consistently underestimate CpG sites in ancestral states and the corresponding trinucleotide mutation rates. The violation of the Jukes-Cantor assumptions results in biased ancestral sequences and dilutes the “true” signal of mutational processes in our data.

Given these concerns, reconstructing ancestral states in an unbiased manner requires using a nucleotide substitution model with trinucleotide or greater complexity. However, these models are extremely computationally intensive. The interdependency of neighboring nucleotides in a trinucleotide mutation model means that computing the probability of transition for a single nucleotide requires consideration of all sites in a genomic window. This means that the computational complexity of a nucleotide substitution model scales non-linearly with context size and sequence length. While a single-nucleotide substitution model took roughly one minute to converge to a solution for a given set of sequences, a trinucleotide substitution model took over eight days on the same data ^42^. The large scale of this analyses, which includes almost 40 billion base pairs in alignments across 15 species, means that a model must be very computationally efficient in order to be feasible for us to implement. A few bioinformatics packages are available to implement trinucleotide substitution models but are too computationally intensive for the scale of our analysis ^42,43^.

### C.0.2 Solution: Mutation Rate Estimation with PhyloFit

In response to these shortcomings, we build a new approach for this thesis. We believe that our method of mutation rate estimation accounts for mutation rate heterogeneity and accounts for ancestral states in an unbiased manner. Instead of counting branch-specific mutations from inferred ancestral states, we directly estimate trinucleotide mutation rates. We estimate these parameters with the expectation of Poisson-distributed single-nucleotide mutations.

Since we are only interested in Single Base Substitutions (SBSs), we split our cleaned, aligned MAF files into 16 context-specific MAF files. Each of these context-specific MAF files, which we will call “context subfiles”, contains all sites in the alignment file with specific surrounding nucleotides. For example, the “A-N-G context subfile” contains all nucleotides (N) in the full alignment with an Adenine (A) adjacent to it in the 5’ direction and a Guanine (G) adjacent to it in the 3’ direction.

We then use the PhyloFit, a program in the PHAST genomics package, to estimate the single-nucleotide mutation rates for each context subfile ^60^. We use the unrestricted single nucleotide model to estimate all 12 free parameters of the single nucleotide transition rate matrix for every terminal branch in the alignment using a Continuous Time Markov Chain approach. Let 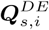 be the generator matrix for species *s* in context subfile *D*-N-*E* in alignment window *i*. We have *D, E* ∈ {A, T, C, G} which represent the fixed surrounding nucleotides for the context subfile. Then, we can express 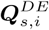 in matrix form as follows:

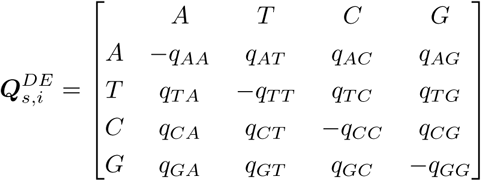

To simplify notation for elements inside the generator matrix, we remove the context-subfile superscripts and species subscript, so 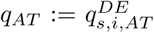. Every off-diagonal element of 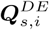 represents a single nucleotide transition rate. Therefore, 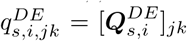 represents the transition rate from nucleotide *j* to nucleotide *k* on the terminal branch for species *s* in the *D*-N-*E* trinucleotide context. In other words, this parameter provides the estimate for the *DjE > DkE* trinucleotide transition rate for species *s* in a given window.

We can interpret this transition rate as the expectation parameter for a Poisson process. Let 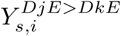 be the total number of *DjE > DkE* mutations on the species *s* branch in the full, non-context specific alignment window *i*. Then,

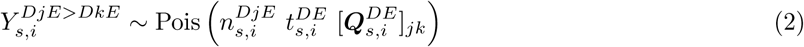

In this notation, 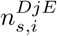 is the number of *DjE* 3-mers in alignment window *i* and 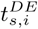 is the PhyloFit-estimated branch length for the *D*-N-*E* context subfile in window *i*. We can interpret 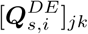 as the trinucleotide transition rate for one nucleotide when 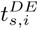 = 1. The subscript *s* reflects the fact that all of these parameters are also species-specific. The number of sites 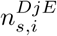 can be trivially extracted from the data, and PhyloFit estimates branch lengths 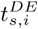 for every context subfile in coordination with the rate parameters 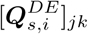 Estimating branch lengths is important in many other areas of phylogenetics, but accurately estimating them requires knowledge of the overall mutation rate for a given genomic region. This is intimately related to the intensity of local mutational processes which we hope to eventually uncover in this analysis. In order to remove PhyloFit’s separation of branch length and generator matrix, we multiply 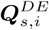 by the scalar branch length 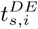. We write 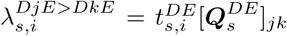. Our species-specific trinucleotide mutation rate matrix across all windows can be written as follows:

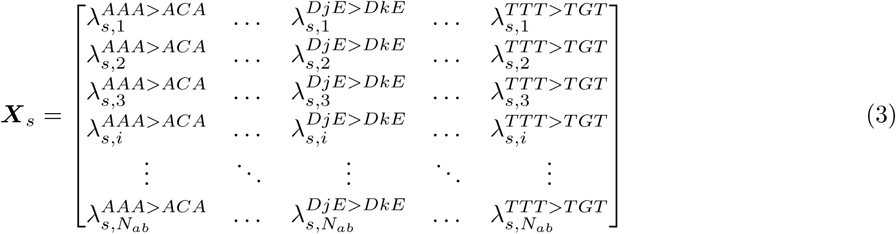

Fixing the trinucleotide context for all base-pairs in the file allows us to interpret the context subfile single nucleotide transition rates as trinucleotide transition rates. In the A-N-G subfile, the estimated C*>*T mutation rate is equivalent to the ACG*>*ATG mutation rate for the whole alignment window. We can estimate any trinucleotide mutation rate using this procedure.

## D Polymorphism data processing

We infer mutational processes for *Papio anubis* (Olive Baboon) and *Pan troglodytes* (Chimpanzees) based on previously published polymorphism data**(author?)** ^50^. Both of these species are included in our analysis of divergence data, and had among the highest numbers of available samples (51 and 29, respectively). Our data processing pipeline was as follows:

**Figure 13:**
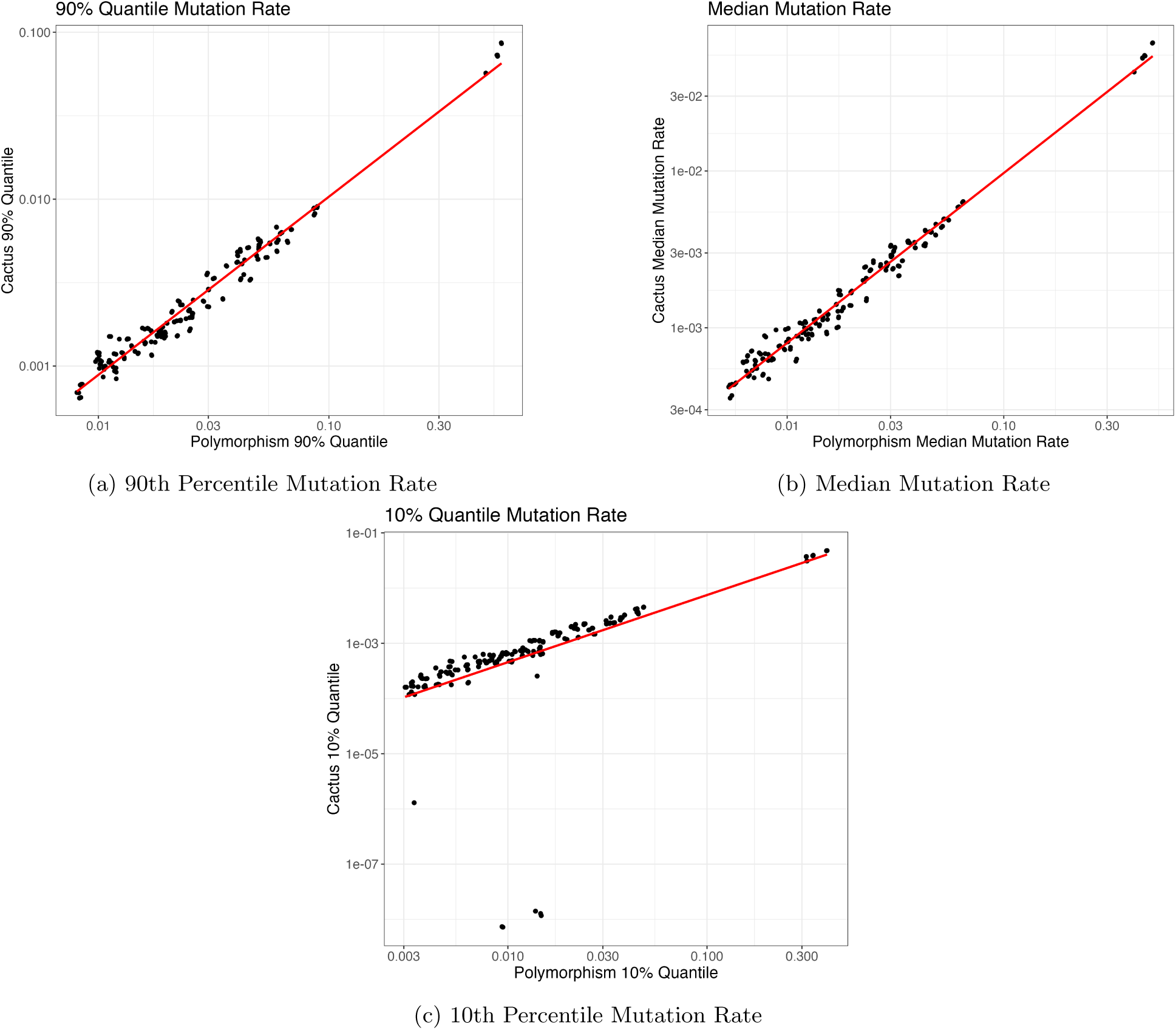
Comparisons of trinucleotide mutation rate distributions between polymorphism (x-axis) and Cactus data (y-axis). Each point on a plot represents a specific trinucleotide mutation type. We compute quantiles for trinucleotide mutation type across all genomic windows. A linear relationship between the two mutation rates, indicated by the red line, indicates consistency in the estimation of the trinucleotide mutation rates.

1. **Merging samples**: Data was provided in compressed Variant Call Format files (vcf.gz) for each sample. We use bcftools merge to combine these different variant calls into one vcf.gz file per species.
2. **Genotyping samples**: From the species-aggregated vcf.gz files, we extracted phase information from all samples at all polymorphic sites using bcftools query. After minimal post-processing, we produce a file with information on reference allele, alternate allele(s), prevalence of each allele in the sample population, and Hardy-Weinberg statistic. This information is calculated individually at every polymorphic site for a given species.
3. **Site filtering**: We excluded sites with a p-value *<* 10*^−^*^5^ with respect to the Hardy-Weinberg statistic. This helps to minimize confounding from selection, population structure, or technical artifacts (i.e. undercalling of heterozygotes during genotyping). Additionally, we removed all sites where the “ancestral” allele had a frequency *<* 0.2 for *Papio anubis*. This is because *Papio anubis* variant calls were performed against the *Macaca mulatta* genome, so sites with low reference allele frequency would likely be the result of divergence from macaques than polymorphism within the baboon population. We considered implementing more aggressive filtering of variants (e.g. only selecting rare polymorphisms), but found that they tended to decrease quality as a result of increased sparsity.
4. **Calculating trinucleotide contexts**: Using the appropriate reference .fasta files from NCBI, we identified the trinucleotide context of every remaining polymorphic site after filtering. Following the approach in ^23^, we calculate the *rate* of trinucletotide mutations in every 1Mb genomic region. These rates are constructed in two steps. First, count the number of trinucleotide mutations of a specific type in a genomic window. This is similar to the mutation counts typically used in somatic analysis. Then, divide the trinucleotide mutation counts by the number of “opportunities” for that mutation type in a genomic window to create a mutation rate. This is done through empirically counting the number of 3-mers of the ancestral state in the genomic window. For example, to calculate the rate of AAA*>*ACA mutations in a population, we would divide the observed number of AAA*>*ACA polymorphisms in a genomic window by the the number of AAA 3-mers in the same genomic window.
5. **Signature extraction**: Steps 1-4 result in the creation of a species-specific trinucleotide rate matrix across 1Mb genomic windows, the same data structure we use in our analysis of divergence-based substitutions (Figure 2). We perform Principal Components Analysis on this data matrix, with the inferred principal components that pass the reflection test representing the extracted mutational processes for a given species.

## E Validation of Inferred Signatures

## F Mutational process clustering

**Figure 14:**
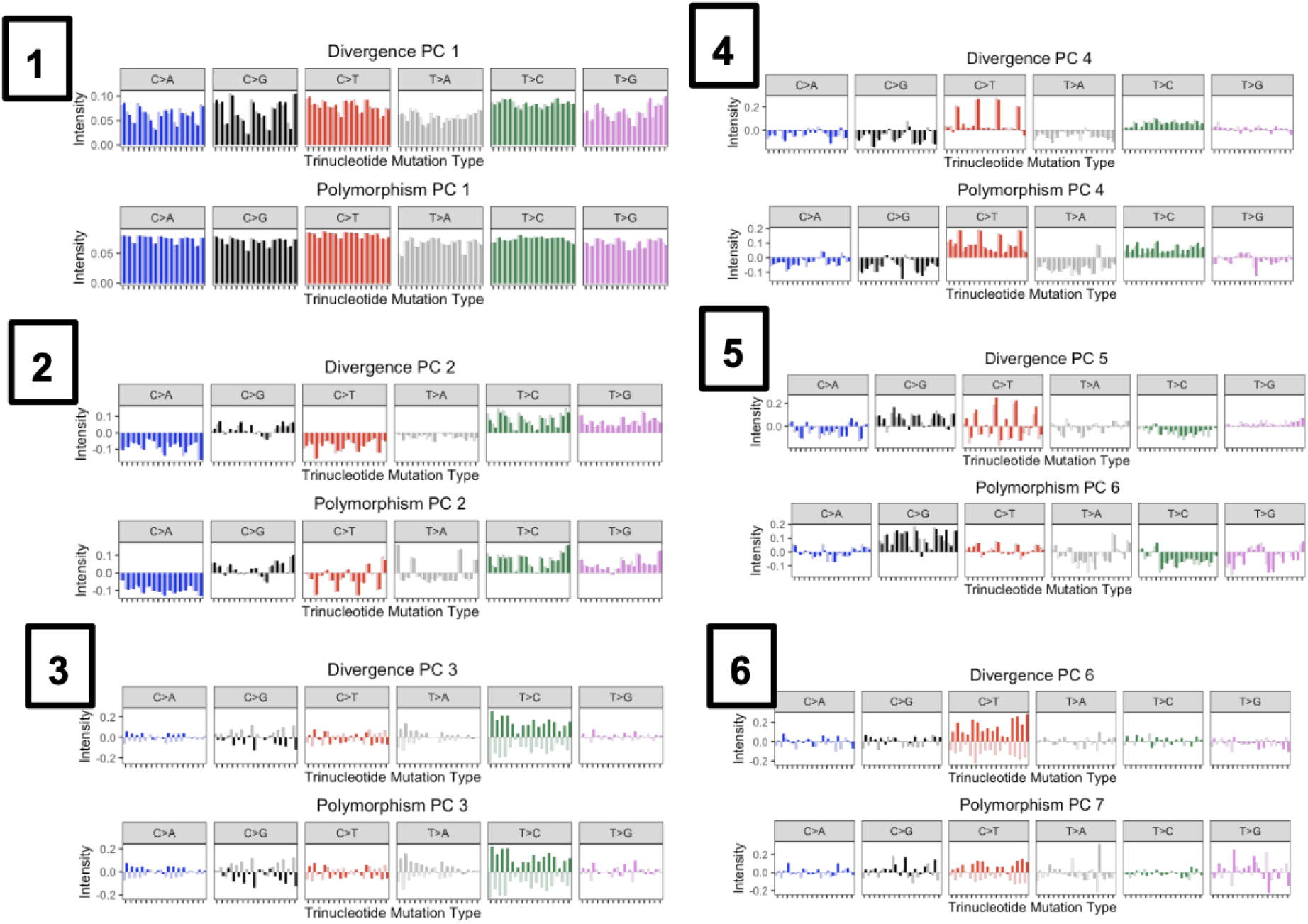
Comparison of divergence-based *Homo sapiens* clade signatures (top panels) and polymorphism-based signatures from *Pan troglodytes* (bottom panels). Boxed numbers indicate the divergence principal components that are being matched.

**Figure 15:**
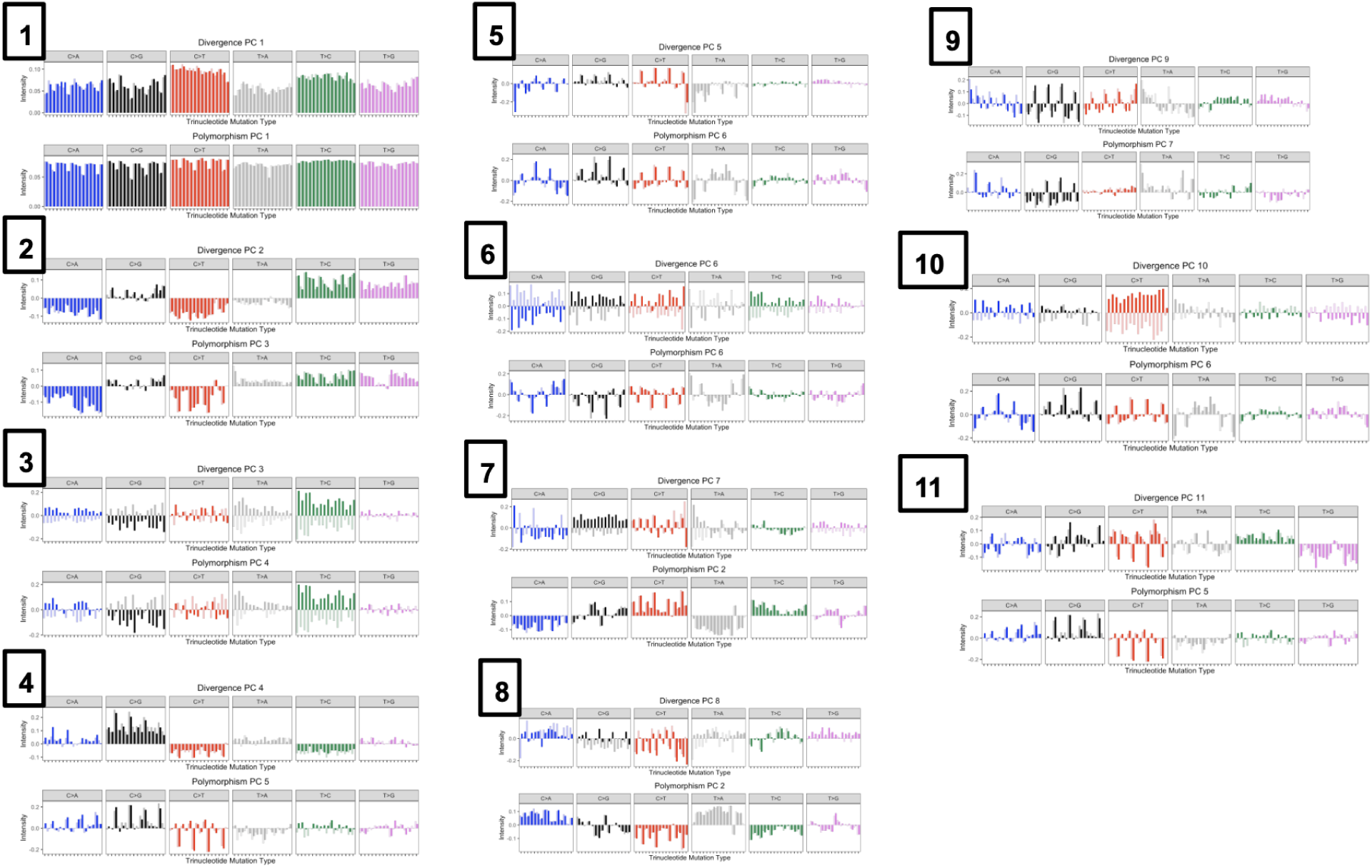
Comparison of divergence-based *Macaca mulatta* clade signatures (top panels) and polymorphism-based signatures from *Papio anubis* (bottom panels). Boxed numbers indicate the divergence principal components that are being matched.

**Figure 16:**
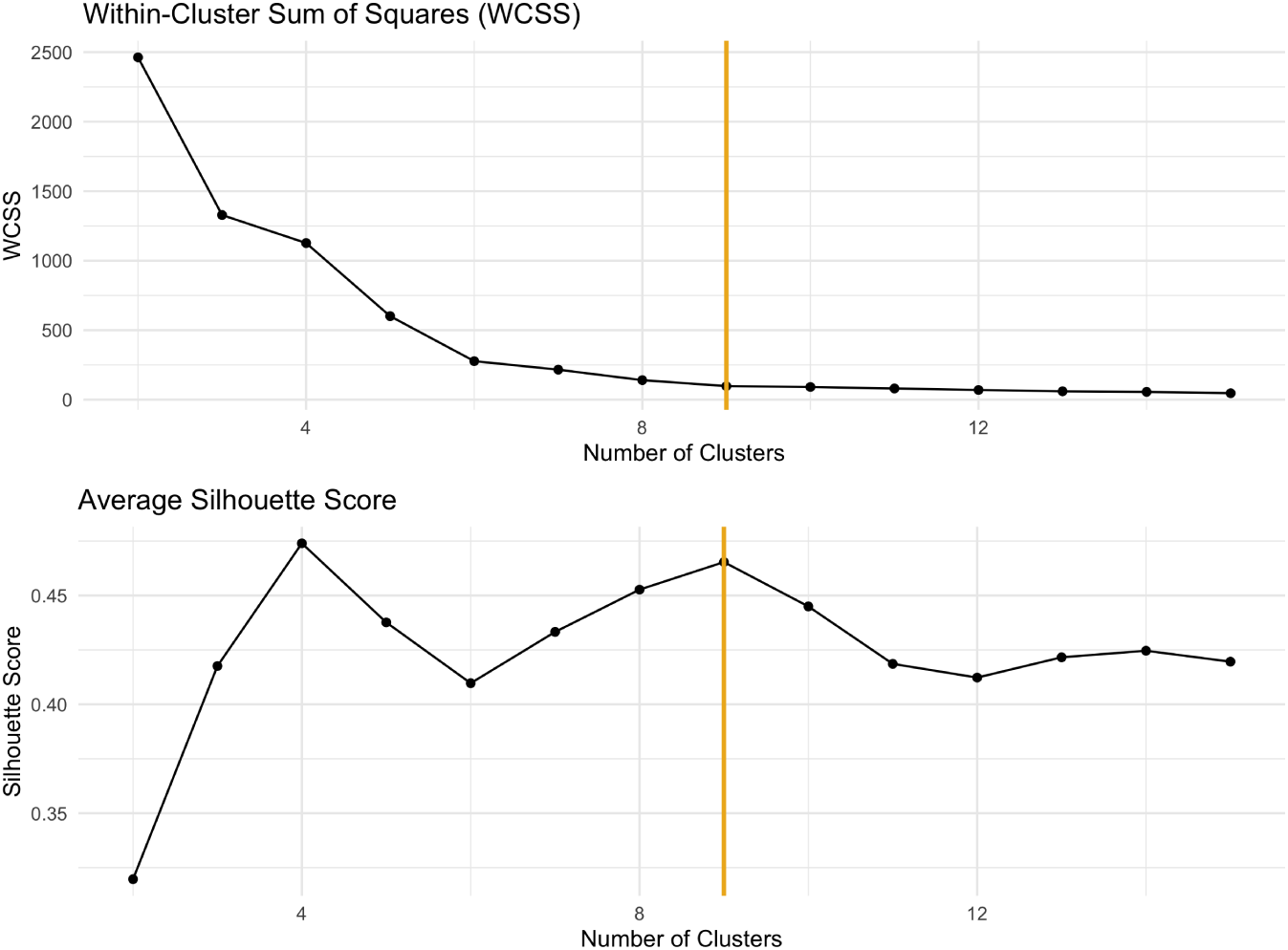
Within-cluster sum of squares (WCSS) for various choices of *k* clusters (top panel), and the average cluster silhouette score ^61^ for choices of *k* (bottom panel).

